# No modular shape changes in the mandible of Andean white-eared opossum (*Didelphis pernigra*, Linnaeus, 1758)

**DOI:** 10.1101/2022.06.09.495434

**Authors:** P.M. Parés-Casanova

## Abstract

Morphological integration and modularity are concepts that refer to the covariation level between the components of a structure. Species of the opossums, genus *Didelphis*, have been the object of several taxonomic and morphometric analyses but no study has so far analysed mandibular morphological integration and modularity at a species-level. The aim of this work was to check whether the body (corpus mandibulae, mandibular corpus) and the ramus (ramus mandibulae, ascending mandibular ramus) are separate modules in *Didelphis pernigra* using a two-dimensional geometric morphometric approach. For this purpose, a sample of hemimandibles from 36 *D. pernigra* (13 males and 23 females) was analysed using 17 landmarks in lateral view. The modularity hypothesis based on different developmental origins was tested, by using the RV coefficient. Later, the integration level was assessed applying a partial least-squares analysis (PLS). The underlying aim was to know whether the traditional division between mandibular body and ramus has a modular basis, as well as the morphological integration level between these two structures. Results reflected that landmarks integration was not uniform throughout the mandible but structured into two distinct modules: ramus and body. Results allow to conclude that allometry plays an important role in shape variation in this species, and that the hypotheses of two-module organization in males cannot be confirmed. Models that accurately represent the biting mechanics will strengthen our understanding of which variables are functionally relevant and how they are relevant to performances, not only masticatories.

## Introduction

Geometric morphometric (GM) relies on multivariate statistics for the quantitative analysis of form, building on more traditional morphometric approaches (Bookstein, 1991) (Mitteroecker & Gunz, 2009) (Adams et al., 2013), allowing to quantify subtle differences in shape that may not be apparent through other means of analyses. Moreover, GM can represent shape variation, since the geometric information encoded in data is preserved throughout the analysis (Rohlf & Bookstein, 1990). GM extracts the information from the data with the Procrustes superimposition method (Mitteroecker & Gunz, 2009) (Rohlf & Bookstein, 1990).

The concept that skeletal form is influenced by extrinsic mechanical forces has been known since a long time, being the role of muscles in the growth and development of skeletal form very complex. Based on this concept, bones would respond to changes in soft tissues (Herring, 1993) (Anderson et al., 2014). The mandibular bone is plastic and so can be modelled during postnatal growth by its interaction with muscles (Anderson et al., 2014).

Morphological integration and modularity are concepts that refer to the covariation level between the components of a structure (Püschel, 2014). Modularity is the property of biological systems to be built of units that are integrated internally and relatively independent from other such units(Klingenberg, 2005). The mandible is an interesting structure for evaluating modularity, especially in a group with a particular mode of reproduction (which impacts the mandible development). The forces of biting are important characteristics of the masticatory apparatus, but few such data are available for the opossum, genus *Didelphis* (Thomason et al., 1990). Species of the opossums have been the object of some taxonomic and morphometric analyses (Thomason et al., 1990) (Astúa, 2015) (Mohamed, 2018) but no study has so far analysed mandibular modular form variation.

Most of the studies on mammal mandible recognize the corpus and the ramus as separate modules (“ascending ramus-alveolar region hypothesis”) (Herring, 1993) (Zelditch et al., 2009) (JojiĆ et al., 2015) (Menegaz & Ravosa, 2017) (Romaniuk, 2018). The aim of this work is to check whether the body (corpus mandibulae, mandibular corpus) and the ramus (ramus mandibulae, ascending mandibular ramus) are separate modules in *Didelphis pernigra*, and if there are sexual differences.

## Material and methods

### Samples

We examined hemimandibles of 36 *Didelphis pernigra* (13 males and 23 females) archived in the collections of the Departamento de Biología of the Universidad del Valle in Cali (Colombia) and Instituto de Ciencias Naturales of the Universidad Nacional de Colombia in Bogotá. Every specimen had been previously identified to the species level and had been collected for other studies. Specimens were caught in the wild from different Colombian localities (not always known). All mandibles were free from any gross skeletal deformities, and none was incomplete. A detailed list of specimens can be obtained upon request to first author.

### Taking photographs and digitizing

Digital images of mandibles were taken with a Nikon D1500 digital camera equipped with an 18-105 mm Nikon DX telephoto lens. Each specimen was placed in the centre of the optical field, with lateral aspect oriented parallel to the image plane. A set of 17 landmarks on the left hemimandible were digitized using TpsDig v.2.16 software. Landmarks 7 to 16 described the body, whereas landmarks 1 to 6 and 17 described the ramus (Table 1 and Figure 1). The landmarks chosen were present on all specimens and were considered to sufficiently summarize the morphology of the lateral aspect of mandible. All images included a ruler for scale. Finally, a generalized full Procrustes fit was performed on two-dimensional landmark coordinates to extract shape information.

**Table 1.** Landmarks (LM) recorded on the analysed morphological mandible. The mandible configuration was divided into subsets of 10 LMs (ascending ramus, LM 7 to 16) and 7 LMs (ramus region, LM 1 to 16, and 17).

|  |  |
| --- | --- |
| 1 | Most dorsal point of *processus coronoideus* |
| 2 | Most dorso-caudal point of *processus coronoideus* |
| 3 | *Incisura mandibulae* |
| 4 | Tip of *processus condylaris* |
| 5 | Orthogonal ventral projection of LM 1 |
| 6 | *Angulus mandibulae* |
| 7 | Most caudal point of molar teeth series |
| 8 | Orthogonal ventral projection of LM 7 |
| 9 | *Caudal foramen mentale* |
| 10 | Orthogonal ventral projection of LM 9 |
| 11 | Orthogonal dorsal projection of LM 12 |
| 12 | *Rostral foramen mentale* |
| 13 | Base of first lower incisor |
| 14 | Most oral point of canine tooth alveolus |
| 15 | Most oral point of first premolar tooth |
| 16 | Limit premolar-molar teeth series |
| 17 | Most dorso-oral point of *processus coronoideus* |

**Figure 1.**
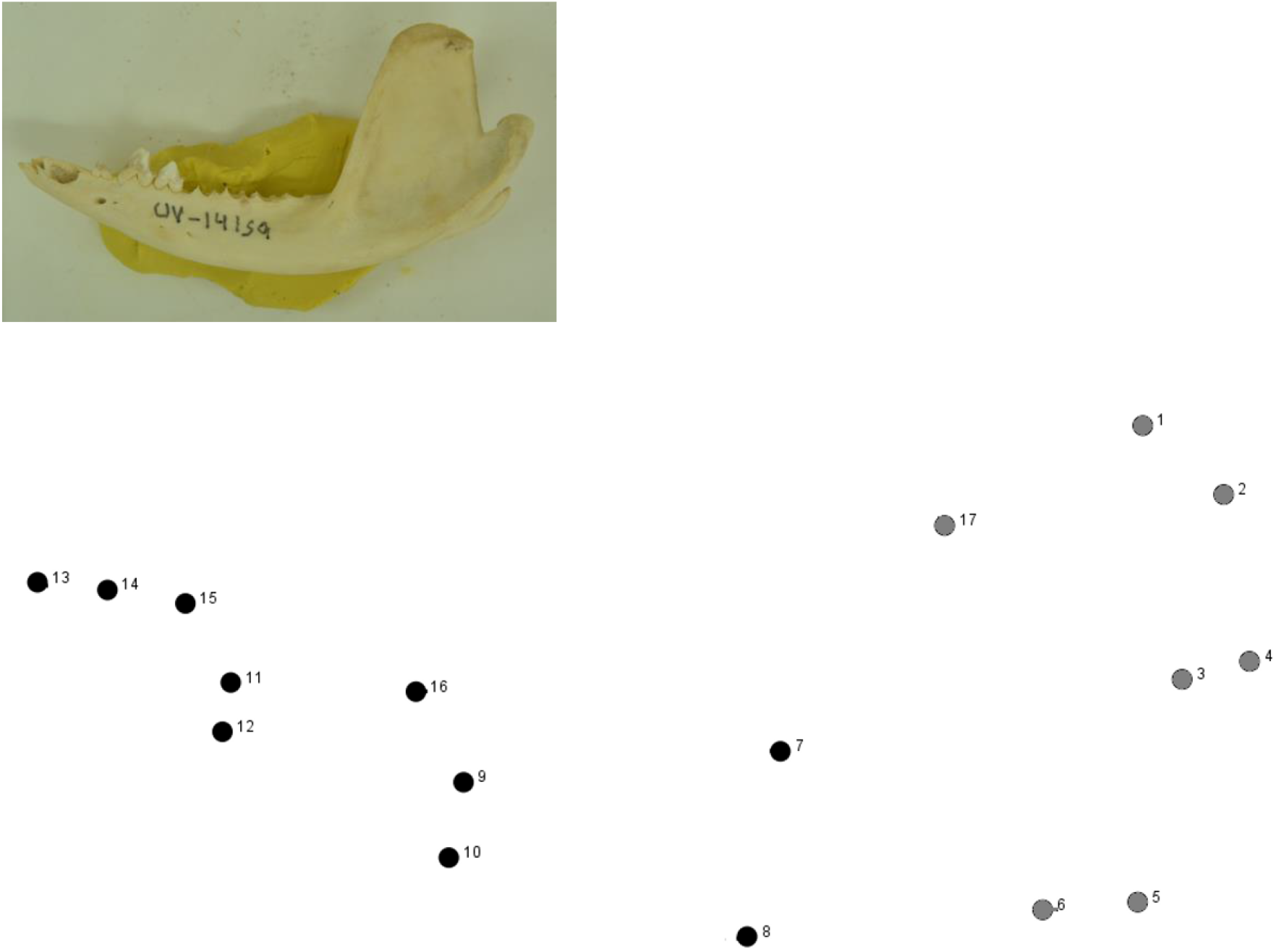
The position of 17 landmarks on the left lateral aspect of hemimandible. Landmarks 7 to 16 described the mandibular corpus (black dots), whereas landmarks 1 to 6 and 17, described the mandibular ascending ramus (grey dots).

### Effect of allometry, integration and modularity

The effects of size can produce global integration throughout the whole landmark configuration and can obscure a possible modular structure (Romaniuk, 2018) so we verified the effect of size correction in *D. pernigra* mandible. The effect of allometry was verified using the multivariate regression of shape (Procrustes coordinates) on size (log10-transformed centroid size) which was treated here as a proxy for general mandible size, and with 10,000 random permutations.

As the two-module hypothesis (subdivision into the alveolar region and the ascending ramus) of data is supported by many authors, we compared subsets of landmarks within those two blocks using RV coefficient as association statistic. The RV coefficient describes the degree of covariation between sets of variables relative to the variation and covariation within sets of variables (Adams, 2016). The proportion of partitions for which the RV coefficient is less than or equal to the RV value for the partition of interest was interpreted as the analogue of a p-value. The hypothesis of modularity is confirmed if the RV coefficient between the hypothesized modules is the lowest or is within the lower tail of the distribution of RVs observed for alternative partitions (JojiĆ et al., 2015). Then, Partial Least Squares (PLS, which is similar to a Principal Component Analysis but using a linear regression model) with 250 rounds reduced the number of variables being observed so patterns were more easily observed in the data. Finally, to assess levels and patterns of shape variations, regression residuals were submitted to a Principal Component Analysis (PCA). All analyses were performed with MorphoJ v.1.06c (Klingenberg, 2011).

## Results

### Allometry

The multivariate regression of the Procrustes coordinates on log10-transformed centroid size showed that allometry was statistically significant (p<.0001) so there was no significative relationship between mandibular shape and size. Log10-transformed centroid size accounted for 13.43% of the total shape variance. Shape changes associated with allometry for both sexes are shown in figure 2, the relationship between mandibular shape and size being quite clear.

**Figure 2.**
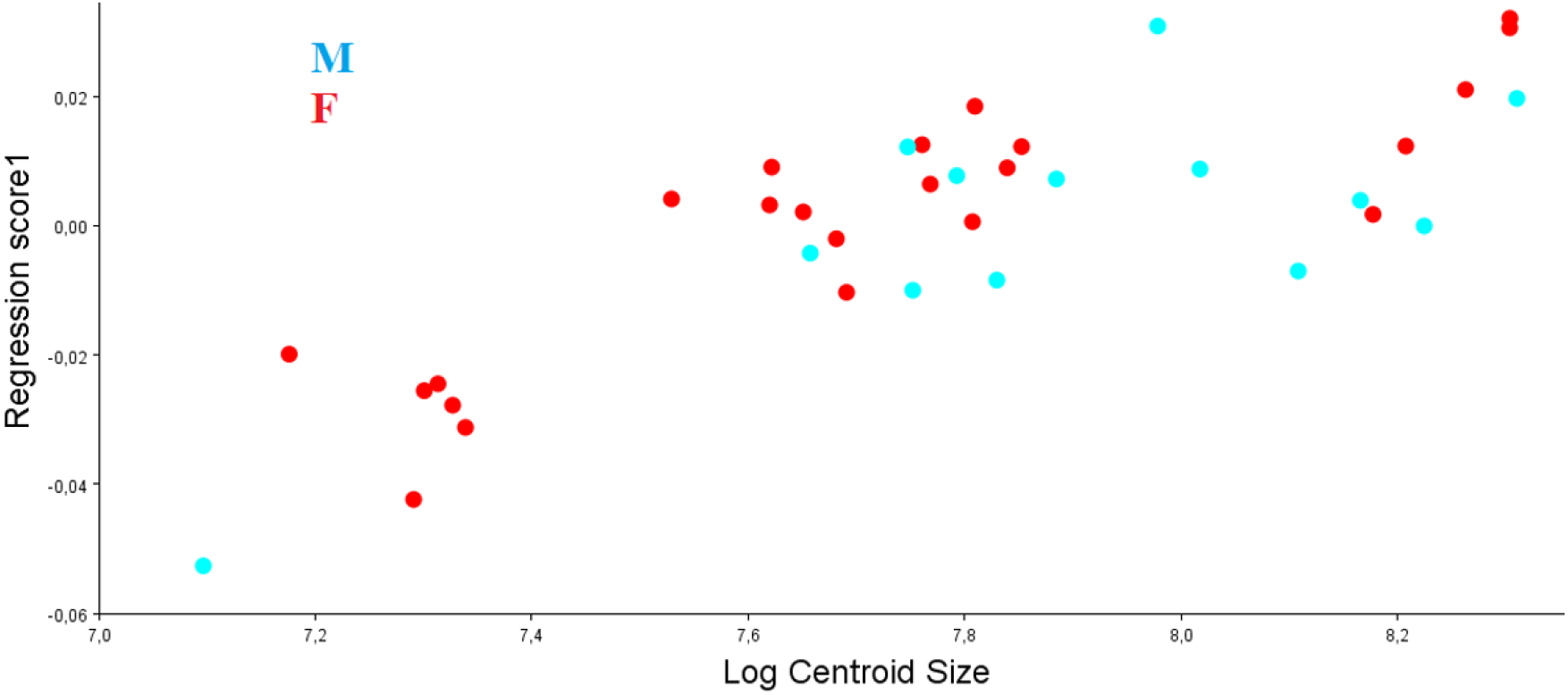
Distribution of *Didelphis pernigra* hemimandibles (13 males and 23 females) in the scatterplot of the shape component (Regression score1) vs log10-transformed centroid size. The multivariate regression showed that allometry was statistically significant (p<.0001, permutation test with 10,000 random permutations). Log10-transformed centroid size accounted for 13.43% of the total shape variance, the relationship between mandibular shape and size being quite clear.

### Integration of mandibular corpus and ramus

PLS-within configuration was made for size-adjusted dataset and considering two-modules. For males, first PLS axes (PLS1) accounted for 100% of the total squared covariance between the mandibular corpus and ramus; singular value=0.00053563, p=0.581. For females, and PLS1 accounted for 100% of the total squared covariance between the mandibular corpus and ramus, too; singular value=0.00041669, p=0.304 for females. So hypothesis of covariation was rejected for both genders. In males, maximum scores of PLS1 were associated with ramus (incisura mandibular and mandibular process), for which correlation between mandibular corpus and ramus was not significative (r=0.893, p>0.05), e.g., not deviating from the correlation expected for random two-module partition of landmarks (Figure 3). In females, maximum scores of PLS1 were associated with ramus, too (Figure 4), for which correlation between mandibular corpus and ramus was not significative, either (r= 0.845, p>0.05).

**Figure 3.**
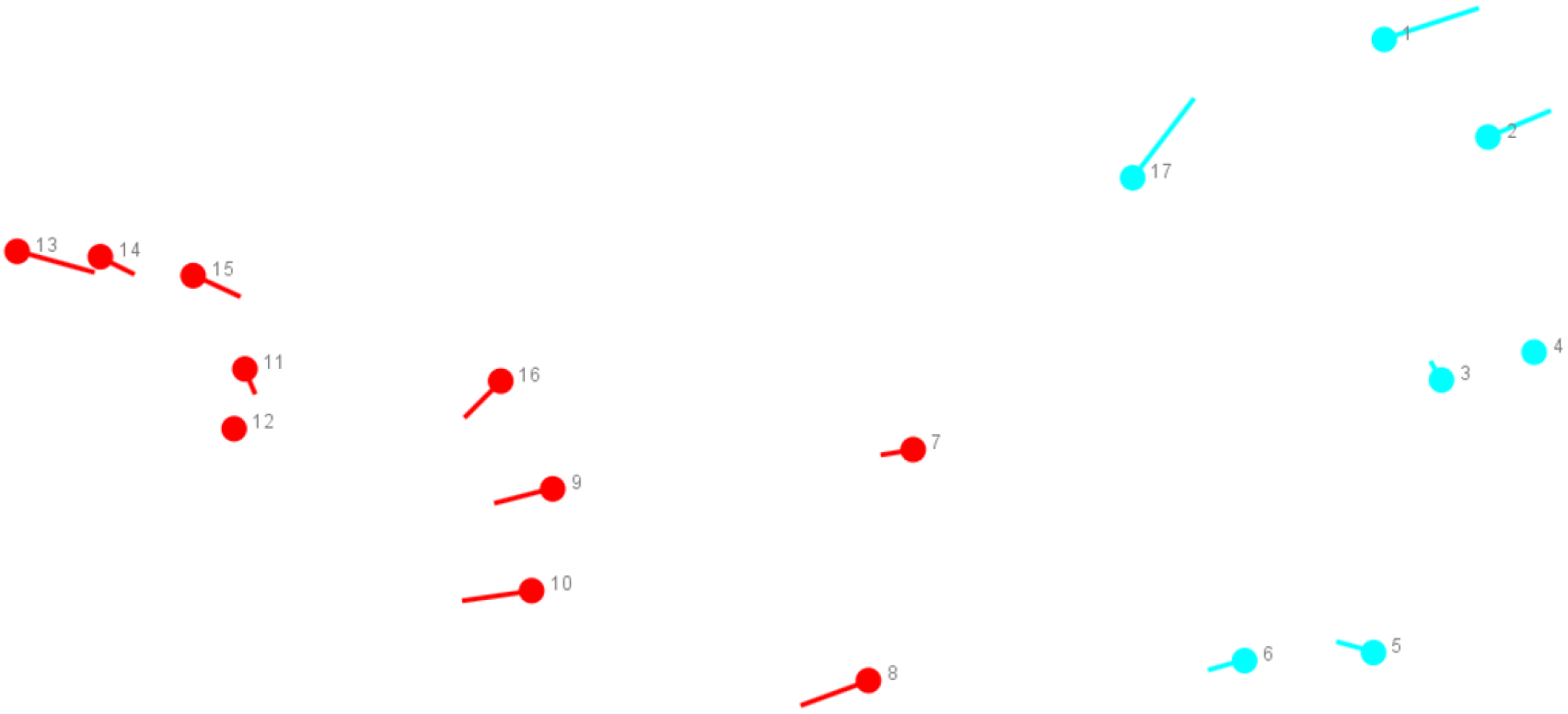
Maximum scores of Partial Least Squares 1 in males, which were associated with ramus (landmarks 1, 2 and 17, located on incisura mandibular and mandibular process).

**Figure 4.**
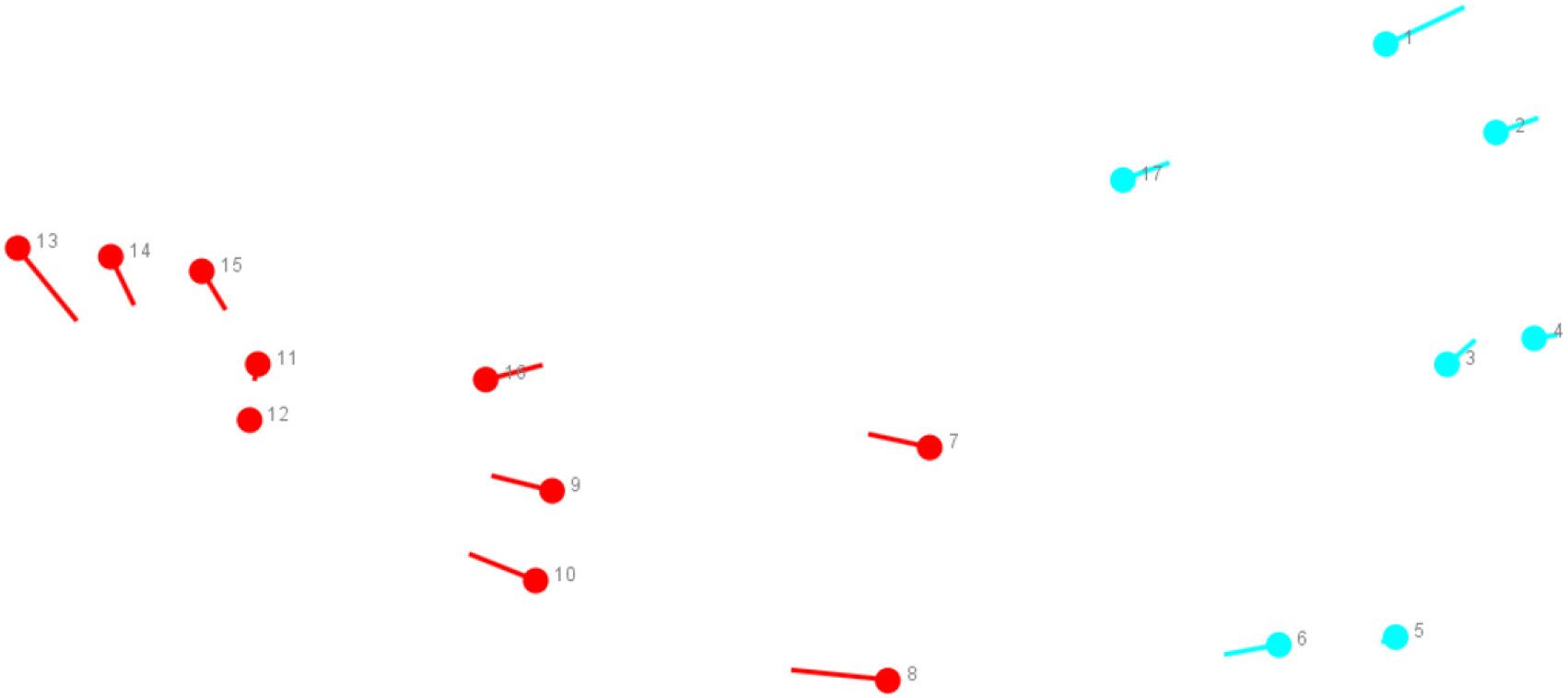
Maximum scores of Partial Least Squares 1 in females, which were associated with ramus (landmarks 1, 2 and 17, located on incisura mandibular and mandibular process).

### Modularity of mandibular corpus and ramus

The hypotheses of two-module organization in males were not confirmed for genders, as 248 out of 1,135 alternative partitions had RVs lower than or equal to the RV coefficient (RV coefficient=0.626, proportion=0.228) calculated for the partition into ramus and body (Figure 5) in males, while for females 135 out of 1,819 alternative partitions had RVs lower than or equal to the RV coefficient (RV coefficient=0.571, proportion=0.074) (Figure 6).

**Figure 5.**
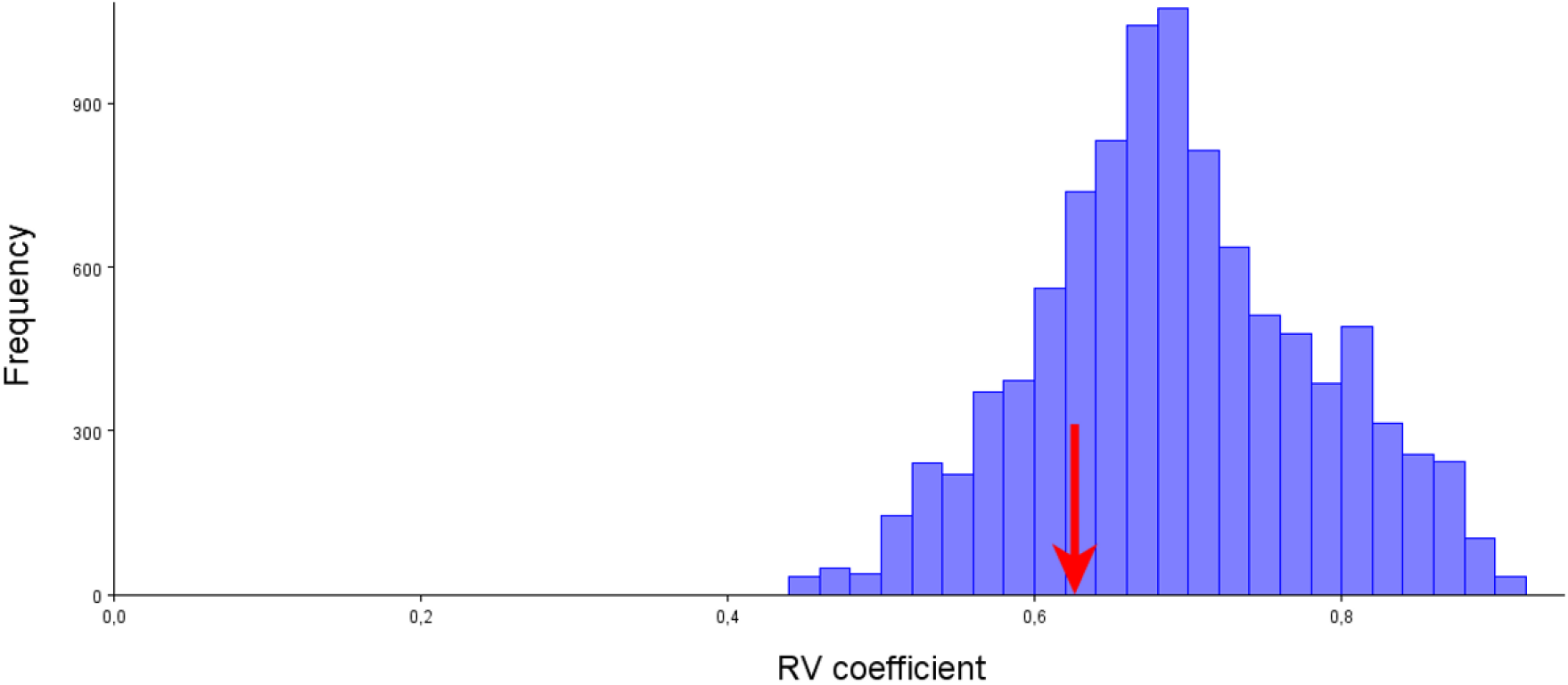
Value of RV coefficient observed for the partition into two hypothesized mandibular modules in males. The hypotheses of two-module organization in males were not confirmed, as 248 out of 1,135 alternative partitions had RVs lower than or equal to the RV coefficient (RV coefficient=0.626, proportion=0.228).

**Figure 6.**
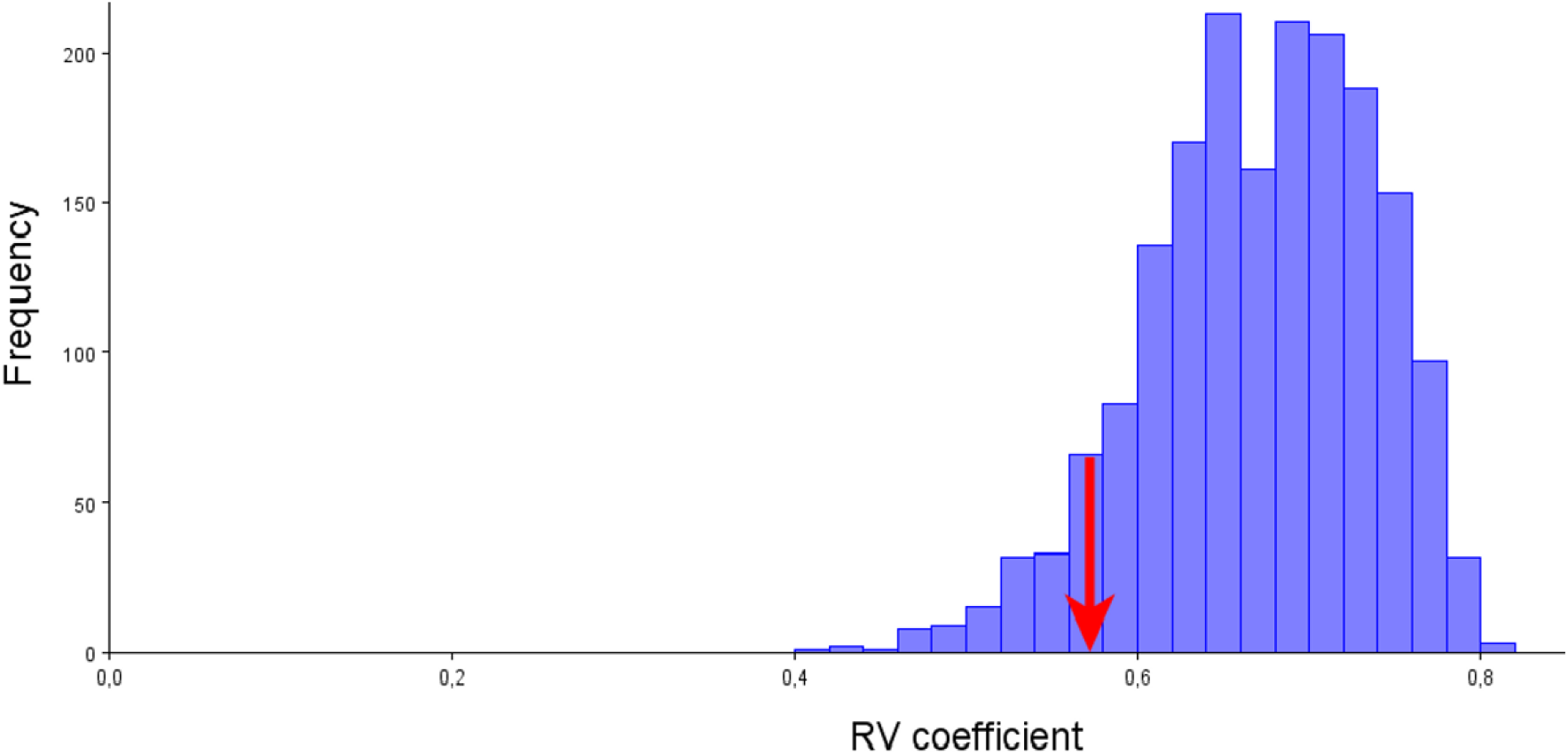
Value of RV coefficient observed for the partition into two hypothesized mandibular modules in females. The hypotheses of two-module organization in males were not confirmed, as 135 out of 1,819 alternative partitions had RVs lower than or equal to the RV coefficient (RV coefficient=0.571, proportion=0.074).

### Principal Component Analysis

The first two PCs explained a low amount of the total shape variation (PC1+PC2 = 33.3%+13.0% = 46.39%). Landmarks had similar loadings.

## Discussion

South American opossums have adapted to a variety of habitats and diets (Cáceres, 2002) (Astúa, 2015). Diet adaptation has a response on mandible, helping animals to feed more effectively (Menegaz & Ravosa, 2017). Mechanical advantage (e.g., bite force) implies therefore more efficient conversion of masticatory muscle force to bite force. In vertebrates, bite force is an important performance trait that can be linked to whole-organism performance because it is relevant to several functions that may impact fitness (Davis et al., 2010). Allometry plays an important role in shape variation in *D. pernigra*, as it has been stated across *Didelphis* species (Astúa, 2015).

Morphological modules are those structures that have components that strongly covary, which in turn are relatively independent to other modules (Püschel, 2014). Morphological integration is the coordinated morphological variation of a functional whole (Püschel, 2014). Results points to weak modularity of the *D. pernigra* mandible. The shape changes are associated with the non-separate (non-discrete) parts of mandible. Traits of shape variation which are specific to these parts make up a low proportion of the total variation as are reflected by the first two principal components.

Future studies on mandibular integration and modularity at multiple levels of variation may shed more light on these important features of morphological variability in other species.

## Acknowledgements

I thank the institutions and people giving us access to their collections: Catalina Cárdenas and Hugo Fernando López for access to the Instituto de Ciencias Naturales of the Universidad Nacional de Colombia collection, and Óscar Murillo, to the Departamento de Biología of the Universidad del Valle collection, and anonymous reviewers for thorough and useful comments that helped us improved the manuscript.

## Conflicts of Interest

The author declares no conflicts of interest.

## Notes

### Competing Interest Statement

The authors have declared no competing interest.

